# Tracing Cellular Senescence in Bone: Time-Dependent Changes in Osteocyte Cytoskeleton Mechanics and Morphology

**DOI:** 10.1101/2024.09.28.615585

**Authors:** Maryam Tilton, Junhan Liao, Chanul Kim, Hossein Shaygani, Maria Astudillo Potes, Domenic Cordova, James L. Kirkland, Kyle M. Miller

## Abstract

Aging-related bone loss significantly impacts the growing elderly population globally, leading to debilitating conditions such as osteoporosis. Senescent osteocytes play a crucial role in the aging process of bone. This longitudinal study examines the impact of continuous local and paracrine exposure to senescence-associated secretory phenotype (SASP) factors on senescence-associated biophysical and biomolecular markers in osteocytes. We found significant cytoskeletal stiffening in irradiated osteocytes, accompanied by expansion of F-actin areas and a decline in dendritic integrity. These changes, correlating with alterations in pro-inflammatory cytokine levels and osteocyte-specific gene expression, support the reliability of biophysical markers for identifying senescent osteocytes. Notably, local accumulation of SASP factors had a more pronounced impact on osteocyte properties than paracrine effects, suggesting that the interplay between local and paracrine exposure could substantially influence cellular aging. This study underscores the importance of osteocyte mechanical and morphological properties as biophysical markers of senescence, highlighting their time-dependence and differential effects of local and paracrine SASP exposure. Collectively, our investigation into biophysical senescence markers offer unique and reliable functional hallmarks for non-invasive identification of senescent osteocytes, providing insights that could inform therapeutic strategies to mitigate aging-related bone loss.

## 1. Introduction

Age-related bone loss impacts over 43 million Americans aged 50 and older, with a projected 47.5% increase in prevalence among those aged 65 and over. [1] This bone loss contributes to osteoporosis and fragility fractures, which remain leading causes of disability, morbidity, and mortality among the elderly. With the global demographic shift towards an aging population, where over 11,000 Americans turn 50 daily and projections indicate that by 2030, all “baby boomers” will be at least 65 years old, [2–4] the urgency to understand the degenerative mechanisms of bone loss becomes a public health priority.

Bone loss is associated with the increased accumulation of senescent cells (SnCs), including senescent osteocytes, and their senescent-associated secretory phenotypes (SASP).[5–8] Indeed, most cells undergoing senescence develop SASPs that include factors with proinflammatory, proapoptotic, and pro-fibrotic effects on neighboring cells.[9] Accumulation of SnCs can have local and systemic detrimental effects due in part to SASP factors.[9,10] To a certain extent, SnCs can be cleared naturally by innate and adaptive immune cells. However, SnCs expressing pro-inflammatory and proapoptotic SASP factors can induce paracrine and endocrine signals, spreading senescence at a rate exceeding immune clearance of existing or newly developed SnCs. This results in accumulation of SnCs at different pathological sites with age. [11] Specifically, SASP factors can induce senescence in normal, non-SnCs both locally and systemically. [12]

SASP factors include pro-inflammatory cytokines, chemokines, growth factors, proteases, and bio-active lipids. [13] In bone, the accumulation of SnCs disrupts the balance between bone formation and resorption, driving the progression of osteoporosis and increasing the risk of fractures. [14,15] The SASP factors secreted by these cells alter the bone microenvironment, enhancing osteoclastic activity and reducing bone density.

The cytoskeleton of cells consists of actin microfilaments, intermediate filaments, and microtubules, providing mechanical support, maintaining cell shape, and playing critical roles in intracellular transport, cell division, and molecular signaling.[16,17] Under mechanical stimulation, the cytoskeleton remodels, including the reorganization of stress fibers, which supports cellular functions by adapting to changes in their mechanical environment.[18–20] Osteocytes, the most abundant skeletal cells, are master regulators of bone homeostasis, being endocrine cells that regulate phosphate metabolism in organs such as the kidney and parathyroid, and most importantly, acting as strain gauges, regulating bone mechanosensation and mechanotransduction.[21–24] Specifically, the osteocyte cytoskeleton is crucial to their function, including cell-extracellular matrix (ECM) interactions via focal adhesions [25,26] Previous studies have demonstrated that SnCs such as senescent mesenchymal stem cells (MSCs) and epithelial cells, undergo alterations in cytoskeletal architecture, potentially impairing their mechanotransduction capabilities. These cytoskeletal changes in aging osteocytes offer opportunities to investigate unique senescence-related biophysical markers, reflecting alterations in their mechanobiological functions.

Cellular senescence is characterized by essentially irreversible proliferative arrest, altered chromatin organization, apoptosis resistance, tumor suppressor activation, oncogenic mutation, and increased protein synthesis.[27–31] Cellular senescence, a state of irreversible cell-cycle arrest, can be induced by various intrinsic (e.g. DNA damage, telomere shortening or dysfunction, oncogene activation, loss of tumor suppressor functions) and extrinsic (e.g. ultraviolet radiation and chemotherapeutic agents) pro-senescence stressors.[32,33] Among these, nuclear DNA damage is a principal trigger of senescence, as it activates the p53-p21 pathway to halt cell proliferation, while epigenetic changes drive senescence via the p16-RB pathway. While senescence is primarily associated with the activation of tumor suppressors, certain oncogenic pathways have been shown to trigger senescence under specific conditions, further reflecting the complexity of the process.[34–36] These processes are highly context-dependent, with variability in the stimuli leading to senescence across different cell types, tissues, and even within heterogeneous cell populations.[37,38] This variability complicates the identification of a universal marker for SnCs. For instance, markers such as Cdkn2a/p16^Ink4a^, which reflects tumor suppressor pathway activation, and Cdkn1a/p21^Cip1^, associated with cell cycle arrest, show variable expression levels in different cellular contexts.[39–42] Senescence associated β-Galactosidase (SA-β-Gal) activity, while commonly used,[43–45] is neither exclusive to SnCs nor consistent across cell populations such as MSCs and epithelial cells. [38,46–48] Similarly, chromatin rearrangements and the formation of senescence-associated heterochromatin foci (SAHF) are observable in some but not all SnC types.[38,49] Recent reviews emphasize the necessity of combining multiple markers, such as transcriptional signatures (e.g., SASP-related gene panels) with biophysical and functional assays, to increase diagnostic accuracy. [37,38,50,51] This multi-marker approach is crucial given the sexual dimorphism in marker expression reported in human and animal models.

Commonly used pro-senescence stressors to study senescence include ionizing radiation, oxidative stress induced by reactive oxygen species (ROS), and chemotherapeutic agents such as doxorubicin.[32,33] These stressors are widely known to induce SASP-related phenotypes, characterized by the release of cytokines, chemokines, growth factors, and proteases that influence local and systemic environments.[51–54] Among these stressors, ionizing radiation is particularly well-suited for in vitro senescence models due to its ability to cause extensive DNA damage, triggering the DNA damage response (DDR) and activating senescence pathways with high reproducibility.[7,38,55–57] Previous studies, including our own, have demonstrated that irradiation effectively models senescence in osteocytes, inducing markers such as *Cdkn2a* (*p16Ink4a*) and *Cdkn1a* (*p21Cip1*), increasing SA-β-Gal activity, and impairing mechanobiological properties.[56,57] These characteristics, combined with precise dosing control, highlight the suitability of irradiation for investigating senescence-associated biophysical changes in osteocytes.

Osteocyte mechanobiology is fundamentally linked to cytoskeletal mechanics and morphology, which are essential for maintaining bone health and function. Alterations in cell mechanics and morphology are not exclusively indicative of senescence but can signal various cellular states or responses. In this study, we specifically probe these changes in the context of exposure to pro-senescence stressors (*e.g*., irradiation) in bone cells. By examining these alterations when cells face such stressors, we aim to establish a novel set of biophysical fingerprints that uniquely characterize cellular senescence within the bone microenvironment. We specifically explore the effects of local SASP accumulation over time on these markers, as well as the paracrine influences of SASP factors through conditioned medium (CM) experiments. By supplying CM to non-irradiated osteocyte cultures, both immediately and after generating a mature healthy culture, we aim to mimic the exposure of non-senescent cells to a SASP-rich environment and investigate their responses. Our findings highlight alterations in cytoskeletal architecture and mechanics correlated to the local and paracrine effects of pro-senescence stressors, which potentially could reflect key osteocyte mechanobiological functions. This approach not only enhances our understanding of the direct and indirect effects of SASP factors but also provides insight into the aging process of bone through the lens of osteocyte biophysical changes.

## 2. Methods

### 2.1 Cell culture and *in vitro* senescence induction

To focus on senescence-associated changes in osteocyte biophysical properties, we employed immortalized osteocytes, specifically MLO-Y4 cells. The mouse-derived MLO-Y4 cells were sourced from Prof. Linda Bonewald’s lab at Indiana University. This approach mitigated the variabilities introduced by heterogeneous primary bone cell cultures. Osteocytes were cultured following established protocols. [58] Briefly, cells were passaged twice before being plated at a density of 2 ×10^4^ cells/35-mm^2^ well. The bottom of the well plates was coated with rat tail collagen type I (0.5 mg/mL, Sigma-Aldrich) and filter-sterilized 0.02 M acetic acid (diluted from glacial, ReagentPlus®, ≥99%, Sigma-Aldrich). The growth media consisted of α-Minimum Essential Medium Eagle (αMEM, Sigma-Aldrich), supplemented with 5% fetal bovine serum (FBS, Gibco), 5% calf serum (CS, Sigma-Aldrich), [59,60] and 1% penicillin-streptomycin (P/S, 10,000 U/mL, Gibco). Half of the medium volume was replenished every four days. The cells were incubated at 37°C in a 5% CO_2_ humidified chamber at confluency of less than 70% to minimize disturbance of cell morphology.

Senescence was induced *in vitro* by a single dose of 10 Gy irradiation (CellRad, Precision X-Ray, *Inc*.). Irradiation was chosen as a method for inducing senescence due to its ability to rapidly activate the DDR, leading to permanent cell cycle arrest. This approach aligns with widely accepted protocols in cellular senescence research.[7,38,55–57] The reproducibility of this method, combined with its precise dosing control, ensures consistent senescence induction across experimental groups.[33,38,50–52] Furthermore, irradiation has been widely used to model osteocyte senescence in vitro, including in MLO-Y4 cells[61] and primary osteocytes[56,57], where it induces markers of senescence such as *Cdkn2a* (*p16*) and *Cdkn1a* (*p21*), increases SA-β-Gal activity, and impairs cellular function. These features make irradiation an effective and clinically relevant model for investigating the impact of senescence on osteocyte mechanobiology. Here, post-irradiation, cells were assessed for biophysical and biomolecular properties on days 7, 14, and 20 (irradiated study groups: D7, D14, and D20). Non-irradiated cells continuously cultured in fresh media served as the Control group (Ctrl).

### 2.2 CM-supplemented groups

To investigate the systemic effects of SASP factors on healthy osteocytes, CM were collected from irradiated osteocyte cultures after 20 days. The CM was centrifuged at 1,500 × g for 5 minutes to remove cellular debris, yielding a consistent and uncontaminated SASP-enriched supernatant. Two groups were established: the immediate CM group (CM-i), where healthy osteocytes were exposed to CM from the start of culture (Day 0), and the delayed CM group (CM-d), where healthy osteocytes were pre-incubated in fresh media for 48 hours before CM supplementation to assess the impact of prior SASP-free growth. Both groups were cultured for 72 hours before biophysical and biomolecular assessments. Cells for imaging and nanoindentation were plated at a density of 1 × 10^5^ cells per 35-mm² well, while those for RT-qPCR were plated at 6 ×10^5^ cells per 35-mm² well, ensuring sufficient material for accurate analysis.

### 2.3 Single-cell nanoindentation

Single-cell micromechanical testing was performed on 50 random osteocytes across two wells using an optical fiber-based interferometry nanoindenter (Pavone, Optics11 Life). Osteocytes were indented with a probe having a 3 µm radius, which is significantly smaller than their surface areas, and with a stiffness of 0.019 N/m. The nanoindentation was performed at a controlled speed of 30 µm/s, reaching a peak load of 0.01 µN, after which the probe retracted at the same speed. The mechanical properties of osteocyte cytoskeleton, characterized by Young’s modulus, were analyzed using Hertzian contact mechanics [62,63]. Indentation curve fitting was conducted using DataViewer V2.5.2 (Optic11 Life) over the linear region on the loading curve, accepting R^2^ > 0.90 (**Figure S2**).

### 2.4 Gene expression

Gene expression analysis was conducted according to the manufacturer’s protocols. RNA was initially extracted from the cells using TRIzol (Invitrogen) followed by purification with the GeneJET RNA Purification Kit (Thermo Fisher Scientific). The concentration of RNA was measured using a NanoDrop Microvolume Spectrophotometer (Thermo Fisher Scientific). Subsequently, cDNA was synthesized from the purified RNA after genomic DNA (gDNA) digestion, using SuperScript™ IV VILO™ Master Mix with ezDNase (Invitrogen). Real-time quantitative Polymerase Chain Reaction (RT-qPCR) was performed using PowerTrack™ SYBR Green Master Mix (Applied Biosystems) with the specific forward and reverse primers listed in **Supplementary Table S1**. The RT-qPCR reactions were conducted on a Quantstudio 6 Pro system (Applied Biosystems). Gene expression results were normalized to Gapdh as the housekeeping gene using the Comparative Ct (2^-ΔΔCt^) method.

### 2.5 SA-**β**-Gal staining

Following the manufacturer’s protocol, cells were fixed and stained with SA-β-Gal (Cell Signaling Technology). After staining, the cells were incubated at 37°C overnight. Cells displaying a blue color were considered to be SA-β-Gal positive. Images were captured using a brightfield microscope equipped with an Olympus XC30 camera, taking five random fields of view per well. The SA-β-Gal activity was quantified by counting the number of positive cells and normalizing this count to the total number of cells observed in each micrograph.

### 2.6 Immunofluorescent (IF) staining

Following the manufacturer’s protocol, cells were fixed in 4% paraformaldehyde (PFA) and stained with Cellmask Orange Actin Tracking Stain (Invitrogen) to visualize filamentous actin (F-actin). The nuclei were counterstained with 4’,6-diamidino-2-phenylindole (DAPI, Invitrogen). Fluorescent images were captured at seven fields of view using an inverted fluorescent microscope (Optics11 Life) equipped with a Nikon 20x objective. Imaging utilized the TRITC and DAPI channels at 545/570 nm and 350/465 nm (excitation/emission wavelengths), respectively. ImageJ 2.14.0 was used for postprocessing. The morphologies of nuclei, F-actin, and dendrites were analyzed and quantified using Cellprofiler 4.2.6 and NeuriteQuant, [64] respectively. The F-actin area was measured through segmentation and quantification of pixels in each fluorescent image (**Figure S4**). While NeuriteQuant was originally developed for neuron analysis, recent studies have demonstrated its application in osteoblasts for the quantification of dendrite lengths,[65] validating its broader functionality for non-neuronal cells. In this study, the software was adapted to ensure reliable quantification of osteocyte dendritic processes. Specific postprocessing parameters were optimized to account for the unique morphology of osteocytes, including adjustments to thresholding and detection settings, to achieve accurate dendrite segmentation and length measurements. Lastly, the multinucleation phenomena within the cell cultures were quantified by counting the number of multinucleated cells and normalizing over the total number of cells in each micrograph.

### 2.7 Statistical analysis

All the reported data underwent the normality and statistical analysis using GraphPad Prism 10. Both one-way and two-way analysis of variance (ANOVA) with Tukey’s multiple comparison tests were employed, each at a 95% confidence interval. One-way ANOVA with Tukey’s multiple comparison tests was utilized for the statistical analysis of SA-β-Gal activity, Young’s moduli, and morphological quantifications. For the RT-qPCR data analyzing gene expression, either One-way or Two-way ANOVA with Tukey’s multiple comparison tests was applied, depending on the experimental design. Statistical significance was considered to be p < 0.05.

## 3. Results and Discussion

### 3.1.1 Irradiated osteocytes exhibited time-dependent changes in biomolecular markers

After irradiation, SA-β-Gal positivity in osteocytes increased over time, which was visualized by increase in cytoplasmic blue staining. The SA-β-Gal-active osteocytes reached close to 100% of the population on Day 20 post-irradiation (p < 0.0001, **Figure 1A**), confirming successful induction of senescence phenotype by low dose irradiation in these cells.

**Figure 1:**
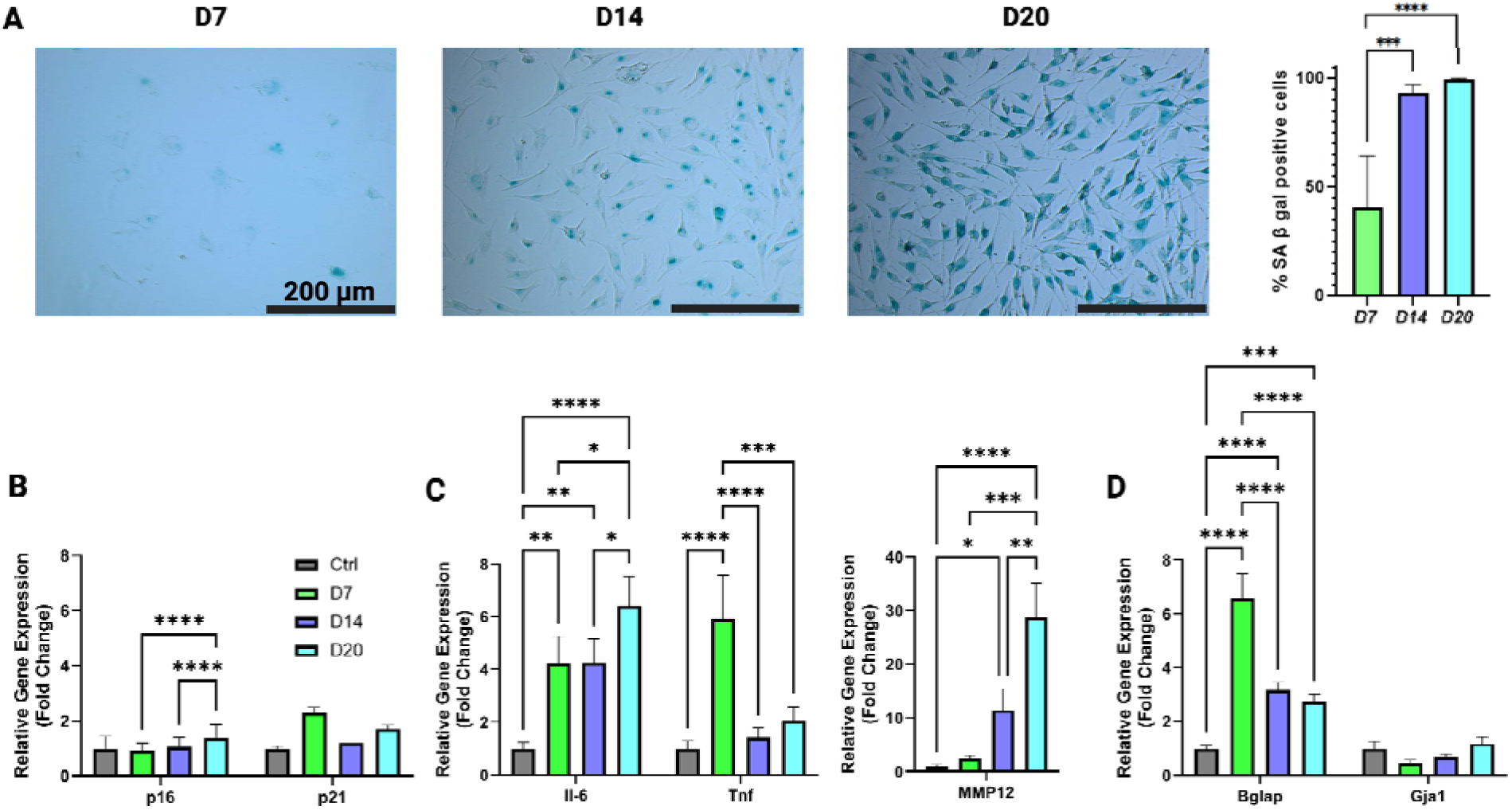
Time-dependent changes in biomolecular markers in irradiated osteocytes driven by local accumulation of SASP factors. **A)** Representative images of SA-β-Gal staining at various time points for irradiated cells (left) with quantification of SA-β-Gal active osteocytes (right). Five images per group were analyzed (n = 5). **B)** RT-qPCR results for senescence, **C)** SASP factors, and **D)** osteocyte markers at all time points. Results are normalized to *Gapdh* housekeeping gene expression and compared to the Control group. Sample size for each group is n = 3, presented as mean ± SD. *: p < 0.05, **: p < 0.01, ***: p < 0.001, ****: p < 0.0001.

Gene expression of senescence markers, SASP factors, and osteocyte markers were measured for all samples. p16^Ink4a^ is a senescence marker that is responsible for regulating the G1/S phase of the cell cycle, [66] while p21^Cip1^ is another senescence marker involved in inhibiting both the G1/S and G2/M phase. [67,68] In the irradiated groups, *p16* and *p21* exhibited gradual increases over time, with *p21* showing an over 2-fold increase compared to the control group by Day 7 (**Figure 1B**). However, these changes were not statistically significant. In contrast, the significant increase in SA-β-Gal positivity across all time points, along with the upregulation of pro-inflammatory SASP markers provides robust evidence for senescence induction in irradiated osteocytes. These findings indicate disruptions of the cell cycle in irradiated osteocytes, which as typical for senescent cells.

The inflammatory response in osteocytes can be reflected by expression levels of interleukin-6 (*Il-6*) and tumor necrosis factor (*Tnf*). The cytokine *Il-6* is a SASP factor that not only stimulates osteoclast differentiation and regulates immune responses but also modulates inflammation. Expression of *Il-6* increased significantly over time in the irradiated groups compared to controls (p < 0.01, **Figure 1C**), suggesting an escalated inflammatory response in irradiated osteocytes. This increase in Il-6 has been previously associated with compromised mechanotransduction in osteocytes [69], linking its accumulation with progressively impaired cellular function [70]. Similarly, expression of Tnf on Day 7 significantly increased, which could potentially signify an increase in receptor activator of nuclear factor κB ligand (Rankl) production and suppression of osteoprotegerin (Opg) [71,72], potentially contributing to bone loss [73]. The elevated expression of both SASP factors in the irradiated groups indicates an increased inflammatory response through autocrine or paracrine effects. The particularly notable time-dependent accumulation of Il-6 underscores its critical role in the senescence-related progression of bone aging.

The ability for osteocytes to regulate bone remodeling was assessed by measuring the levels of matrix metalloproteinase (Mmp12) and bone gamma-carboxyglutamate protein (Bglap). Mmp12, a SASP factor known for its role in ECM degradation, is critical in bone remodeling processes. Its increased activity is associated with accelerated bone resorption and potentially dysfunctional bone remodeling [74]. Expression of Mmp12 was elevated in the Irradiated groups over time, with more than a 20-fold increase by Day 20 compared to the Control (p < 0.0001, **Figure 1C**). This result suggests intensified ECM degradation and possible remodeling dysfunction in these groups.

Osteocalcin (*Bglap*) plays a pivotal role in bone formation and remodeling. [75] During bone formation, *Bglap* enhances matrix mineralization by binding calcium and hydroxyapatite, contributing to the structural integrity of bone.[76,77] In remodeling, *Bglap* deposited in the bone matrix indirectly promotes resorption by stimulating osteoclast activity, thereby coordinating the balance between bone formation and resorption.[76,77] Its dual role in these processes makes it an ideal marker for investigating the metabolic dynamics of bone. In this study, irradiated osteocytes showed significantly increased *Bglap* expression (p < 0.0001 for Day 7 and 14, p < 0.001 for Day 20, Figure 1D), suggesting enhanced bone resorption activity and its potential contribution to overall bone loss during senescence. This finding indicates the potential role of *Bglap* as a key metabolic marker in bone remodeling under senescence-associated conditions.

Gap junction protein alpha 1 (*Gja1*) expression can be linked to osteocyte dendrite function in terms of intercellular communication.[78] Disruption in Gja1 production has been previously associated with age-related osteoclastogenesis and bone loss.[79] In **Figure 1D**, there were no statistically significant changes in Gja1 expression among the groups (Ctrl, D7, D14, and D20). While slight variations were observed, such as a marginal increase in Gja1 expression on Day 20 compared to Day 7, the lack of significant changes in Gja1 expression suggests that intercellular communication via gap junctions may not be severely impaired under the tested conditions.

The gene expression results evaluated by RT-qPCR suggested that over time, the irradiated group exhibited an increase in senescence markers and higher levels of pro-inflammatory cytokines. The increased expression of Bglap further underscored declines in osteocyte mechanobiological function and potential dysregulation of bone resorption.

Importantly, irradiation-induced senescence primarily represents stress-induced premature senescence (SIPS) rather than replicative senescence (RS). [37] However, this method offers distinct advantages, such as rapid senescence induction and precise experimental control.[37] Unlike RS, which arises gradually through telomere attrition, SIPS bypasses telomere erosion and directly activates DDR pathways. [37] It should be noted that cellular responses to ionizing radiation may vary based on cell cycle phase, with proliferating cells being more susceptible to damage than non-dividing cells.[33,37,80] These factors necessitate careful interpretation of findings.

### 3.1.2 Short-term SASP factor exposure induced biomolecular marker changes in healthy osteocytes

Gene expression levels from Day 7 served as the control in order to compare the local and paracrine effects of SASP factors (**Figure 2A**). Expression of *p16* in the CM-i group was similar to that observed on Day 7, while expression in the CM-d group was lower (0.525-fold). Conversely, *p21* expression increased in the CM-i group (1.745-fold), with levels in the CM-d group matching those on Day 7 (see **Figure 2B**). The stable *p16* expression in the CM-i group, compared to the irradiated groups, suggests a disruption in the S phase of the cell cycle, indicating that DNA synthesis in osteocytes exposed to CM-i was significantly impacted by SASP factor exposure. On the other hand, the increase in *p21* levels in the CM-i group suggest potential dysfunction in mitosis, reflecting disruption in the M phase of the cell cycle. Despite the changes in p16 and p21 expressions in CM-d group, this group exhibited minimal SA-β-gal positivity (**Figure S1**).

**Figure 2:**
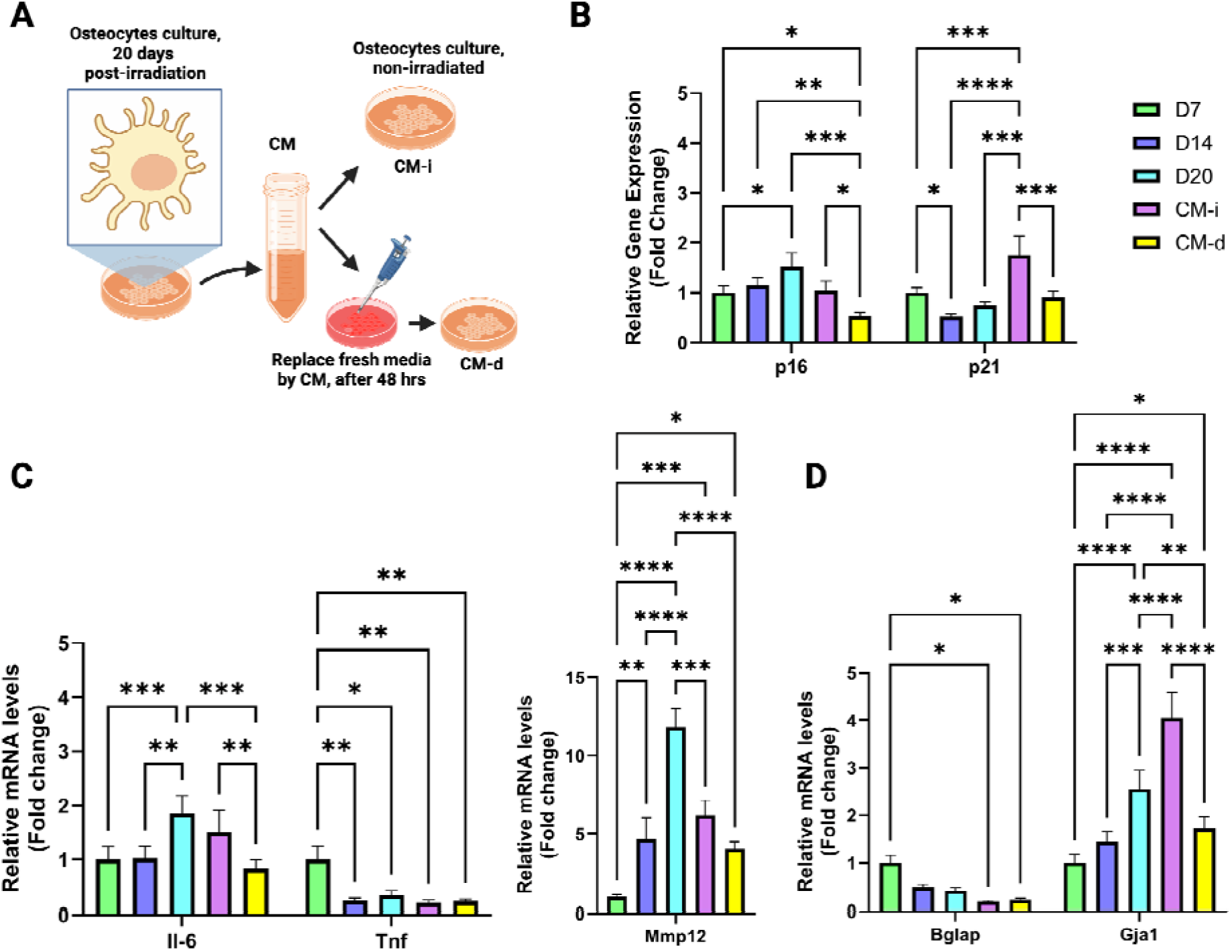
Systemic supplementation with SASP factor-enriched conditioned media induce senescence phenotypes in non-irradiated osteocytes. **A)** Schematic of the CM-i (immediate supplementation) and CM-d (delayed supplementation) culture setups. **B)** RT-qPCR results for senescence, **C)** SASP factors, and **D)** osteocyte markers across all conditions, normalized to *Gapdh* housekeeping gene expression and compared to the Day 7 post-irradiation group. Sample size for each group is n = 3, presented as mean ± SD. *: p < 0.05, **: p < 0.01, ***: p < 0.001, ****: p < 0.0001.

In the CM-d group (**Figure 2A**), *Il6* and *Mmp12* levels were comparable to those observed on Day 7, whereas the CM-i group exhibited a 1.509-fold increase in *Il6*, surpassing the levels noted on Day 14. Conversely, *Tnf* levels in both the CM-i and CM-d groups remained low, aligning more closely with the values observed on Days 14 and 20 rather than Day 7. *Mmp12* levels in both CM-supplemented groups showed a notable increase compared to Day 7 (6.198-fold and 4.047-fold, respectively; see **Figure 2C**). These results suggest that *Il6* and *Mmp12* played a more prominent role in autocrine/paracrine signaling than *Tnf* in the non-irradiated cells within the CM-supplemented groups. Additionally, the immediate incubation of healthy osteocytes in SASP-supplemented CM in the CM-i group may have stimulated a more effective uptake and signaling response to *Il6*, further amplifying its expression. Overall, these findings confirm that healthy osteocytes in both CM-supplemented groups effectively absorbed SASP factors from their conditioned media. Gene expression analysis comparing local versus systemic SASP exposure, normalized to Day 14, further underscores these observations (**Figure S5**). The senescence marker *p21* was significantly increased in both CM-i and CM-d groups compared to Day 14, while *p16* expression remained unchanged and non-significant (**Figure S5-A**). SASP-related cytokines, such as *Il6*, *Mmp12*, and *Tnf*, were comparable to their expression levels on Day 14 (**Figure S5-B and S5-C**). However, *Il6* and *Mmp12* levels in the CM-d group were significantly lower compared to Day 20.

Bglap levels in both CM-supplemented groups were significantly lower than those observed at all local time points, including Day 7, Day 14, and Day 20 (p < 0.05, **Figure 2D**, and **Figure S5-D**). This finding suggests that the paracrine effects of SASP factors were less effective in promoting osteocalcin production compared to their local effects. Similarly, Gja1 levels in both CM-supplemented groups showed an increase from Day 7, with levels in the CM-d group remaining similar to Day 14, whereas CM-i exhibited a significant increase compared to Day 7 (p < 0.0001, **Figure 2D**, and **Figure S5-D**). This increase might be attributed to the abrupt exposure of osteocytes in the CM-i group to SASP factors in the CM triggering a more intensive survival response. Overall, the gene expression results for the CM-supplemented osteocyte confirm effective uptake of SASP factors and suggest potential stimulation of autocrine/paracrine SASP signaling pathways. The distinctly different profiles of senescence and osteocyte markers reflect potential variations in the mechanisms through which paracrine SASP factor exposure influences non-irradiated osteocytes.

### 3.2 Osteocyte morphological changes as biophysical markers for senescence

Various morphological parameters were quantified in our longitudinal study to examine senescence-associated changes (**Figure 3A**). Nuclear blebbing, observed in the irradiated groups (**Figure 3B**), indicated nuclear membrane instability. Additionally, nuclear expansion was noted over time, being particularly significant on Days 14 (p < 0.0001) and 20 (p < 0.001, **Figure 3C**). In contrast, osteocytes in both CM-supplemented groups exhibited no significant changes from the control group, suggesting that short-term exposure to SASP factors may not induce profound alterations in nuclear morphology. Such stability might be attributed to mechanisms such as chromosome unwinding within the nuclei, which potentially help mitigate aging-related DNA damage. The nuclear abnormalities observed in the irradiated groups signify compromised nuclear function in osteocytes, which could lead to phenotypic variations during protein translation as previously observed in other cell types [81,82]. These findings should be further investigated in future studies.

**Figure 3:**
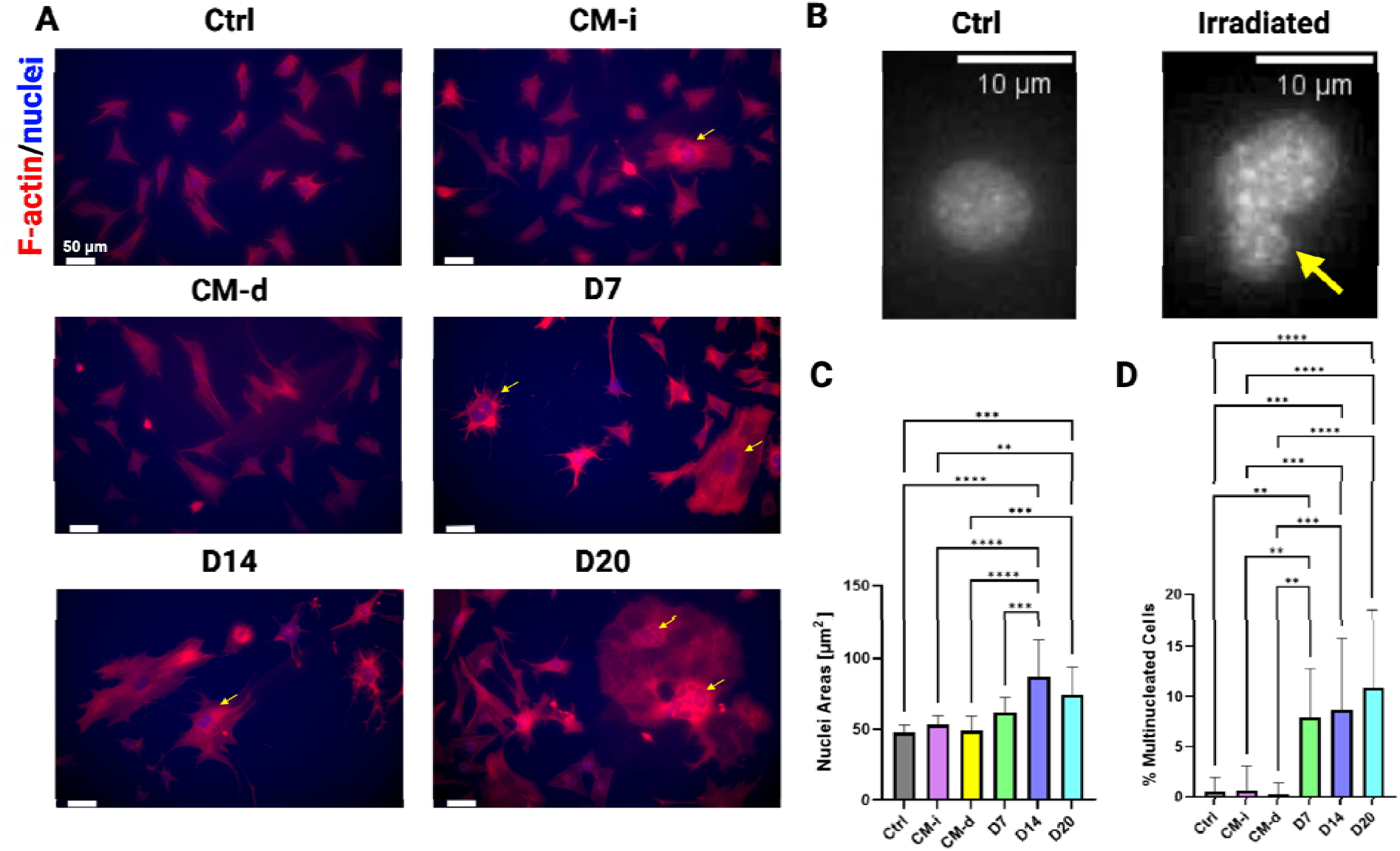
Alterations in F-actin organization and nuclear morphology are associated with local and systemic exposure to SASP factors. **A)** Representative IF micrographs showing F-actin and nuclei staining across all experimental conditions, with yellow arrows highlighting multinucleated cells. Scal bar: 50 µm. **B)** Close-up view of nuclear blebbing in irradiated osteocytes, indicated by a yellow arrow. **C)** Average nuclear areas quantified using CellProfiler from IF micrographs across all conditions. **D)** Percentages of multinucleated osteocytes in each experimental group. Quantitative analysis of osteocyte morphology was performed by examining 14 IF micrographs per group across two cell culture wells (n = 14 for all groups, presented as mean ± SD). Significance levels are denoted as *: p < 0.05, **: p < 0.01, ***: p < 0.001, ****: p < 0.0001.

A significantly elevated population of multinucleated osteocytes was observed in the Irradiated groups, reaching as high as 11% on Day 20 (p < 0.0001, **Figure 3A, 3D**). In contrast, both CM- supplemented groups maintained similarly low levels of multinucleated cells to the Control group. Since senescent osteocytes often exhibit disrupted cell cycles, the observed multinucleation may result from incomplete M phases and/or DNA hyper-replication, similar to findings reported in oncogene-induced senescence studies of primary human fibroblasts [83]. This hypothesis is supported by the increased expression of p21 over time in the Irradiated groups, underscoring the progression of cell cycle disruption. The prevalence of multinucleation in irradiated osteocytes further emphasizes the impact of their local SASP factor accumulation, which appears to be more consequential than short-term SASP exposure from their microenvironment.

F-actin networks in osteocytes from the Irradiated groups exhibited extensive expansion compared to the Control group, characterized by reduced dendricity and non-uniform organization of F-actin structures (**Figures 3A**, **4A**). The actin disassembly (**Figures 3A**, **4A**) and potentially reduced polymerization dynamics are consistent with oxidative stress-induced mechanisms reported in previous studies.[84,85] Specifically, previous studies demonstrated that oxidative stress can alter the actin cytoskeleton through Cys374 oxidation, leading to disruptions in filament stability and polymerization.[84,85] These mechanisms align with our observations and suggest a role for oxidative stress in the observed cytoskeletal changes. However, we acknowledge that direct oxidative stress measurements were not performed in this study. Future experiments will incorporate assays, such as superoxide anion fluorescent probe staining, to directly evaluate reactive oxygen species (ROS) levels and their effects on the osteocyte cytoskeleton.

A progressive time-dependent increase in the average F-actin area was observed in the Irradiated groups (p < 0.0001), significantly exceeding that of the Control and both CM-supplemented groups (**Figure 4B**). This increase could be attributed to abnormal filopodia and lamellipodia formation in senescent cells, which likely contributed to osteocyte flattening and area expansion. The disorganized and expanded F-actin networks may have resulted in spatially heterogeneous mechanical properties within the osteocytes.

**Figure 4:**
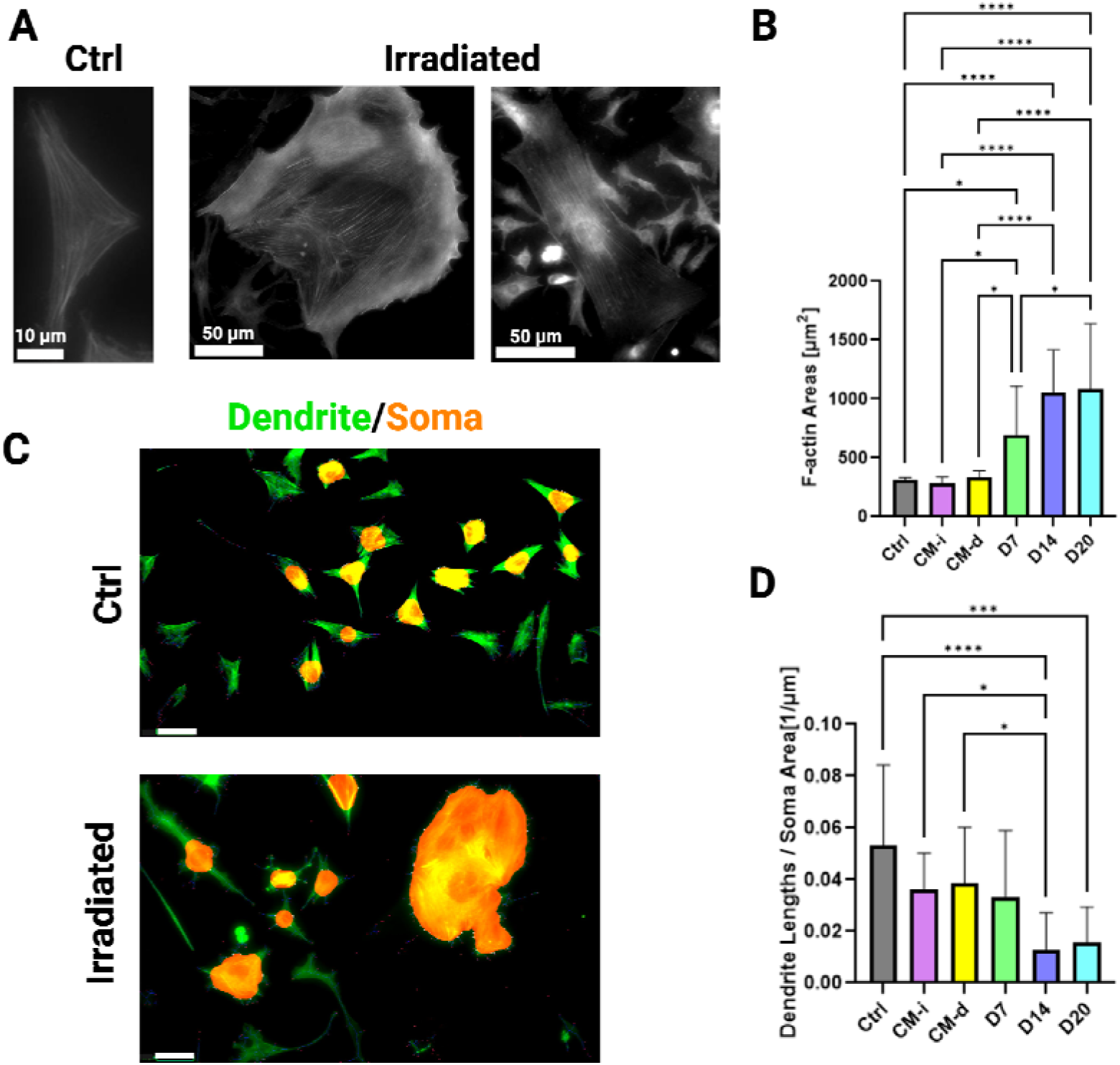
Quantitative analyses of F-actin area and dendritic structures in osteocytes following SASP factor exposure compared to non-senescent controls. **A)** Representative images of F-actin organization in osteocytes from Control and Irradiated groups. Scale bar: Control: 10 µm, Irradiated: 50 µm. **B)** Quantification of average F-actin areas using CellProfiler from IF micrographs across all experimental conditions. **C)** Representative images of semi-automated segmentation and quantification of dendrites and soma performed using NeuriteQuant. Scale bar: 50 µm. **D)** Quantification of average dendrite length normalized to soma area, calculated from IF micrographs of all conditions using NeuriteQuant. All quantitative osteocyte morphology results were obtained by analyzing 14 IF micrographs per group across two cell culture wells (n = 14 for all groups, presented as mean ± SD). Significance levels are indicated as *: p < 0.05, **: p < 0.01, ***: p < 0.001, ****: p < 0.0001.

The temporal accumulation of SASP factors, including *Il6*, *Tnf*, and *Mmp12* (Figure 1C), suggests that SASP-driven signaling may also play a crucial role in inducing oxidative stress and contributing to F-actin expansion. Previous studies have demonstrated that pro-inflammatory cytokines, such as those included in the SASP, can amplify oxidative stress, leading to cytoskeletal remodeling and impaired cellular function.[17,80,86,87] These findings align with interpretations in the field but require further investigation to directly link specific SASP factors to oxidative stress and the biophysical changes observed in osteocytes.

Overall, these findings demonstrate that local SASP factor accumulation significantly altered osteocyte F-actin morphology, potentially impairing their mechanobiological function. In contrast, the paracrine effects of SASP factors were less pronounced in influencing F-actin expansion, likely due to the relatively brief exposure to oxidative stress compared to the longer-term exposure experienced by osteocytes in the Irradiated groups.

The dendrite lengths of irradiated osteocytes were observed to be generally shorter compared to the Control group (see **Figure S3**). Specifically, normalized dendrite lengths in the irradiated groups significantly decreased by Day 14 (p < 0.0001) and continued to decline through Day 20 (p < 0.001, **Figure 4D**). While not statistically significant, dendritic lengths in both CM-supplemented groups were also reduced compared to the Control cells, aligning with measurements from Day 7 (**Figure 4D** and **Figure S3**). To mitigate confounding effects due to variations in F-actin area, dendrite lengths were normalized to soma areas, which was quantified via NeuriteQuant (**Figure 4C**). This analysis underscores the structural alterations induced by both local and systemic SASP effects on osteocytes structure and morphology. In particular, the irradiated osteocytes with local SASP accumulation exhibited diminished dendritic network connectivity, which may suggest reduced intercellular communication and mechanosensation capabilities.

### 3.3 Osteocyte cytoskeleton stiffening as a biophysical marker for senescence

To characterize the longitudinal changes in cytoskeletal stiffness due to local and paracrine exposure to SASP factors, we employed live single-cell nanoindentation (**Figures 5A-B**). At all study time points, osteocytes from the Irradiated groups exhibited significantly higher Young’s modulus (*i.e*. stiffnesses) compared to those in the Control and CM-supplemented groups (see **Figure 5C**). The CM-d group displayed no significant change in stiffness compared to the Control (**Figure 5C**). This increase in cytoskeletal moduli could be linked to the previously demonstrated expansion in F-actin area (see **Figure 4B**).

**Figure 5:**
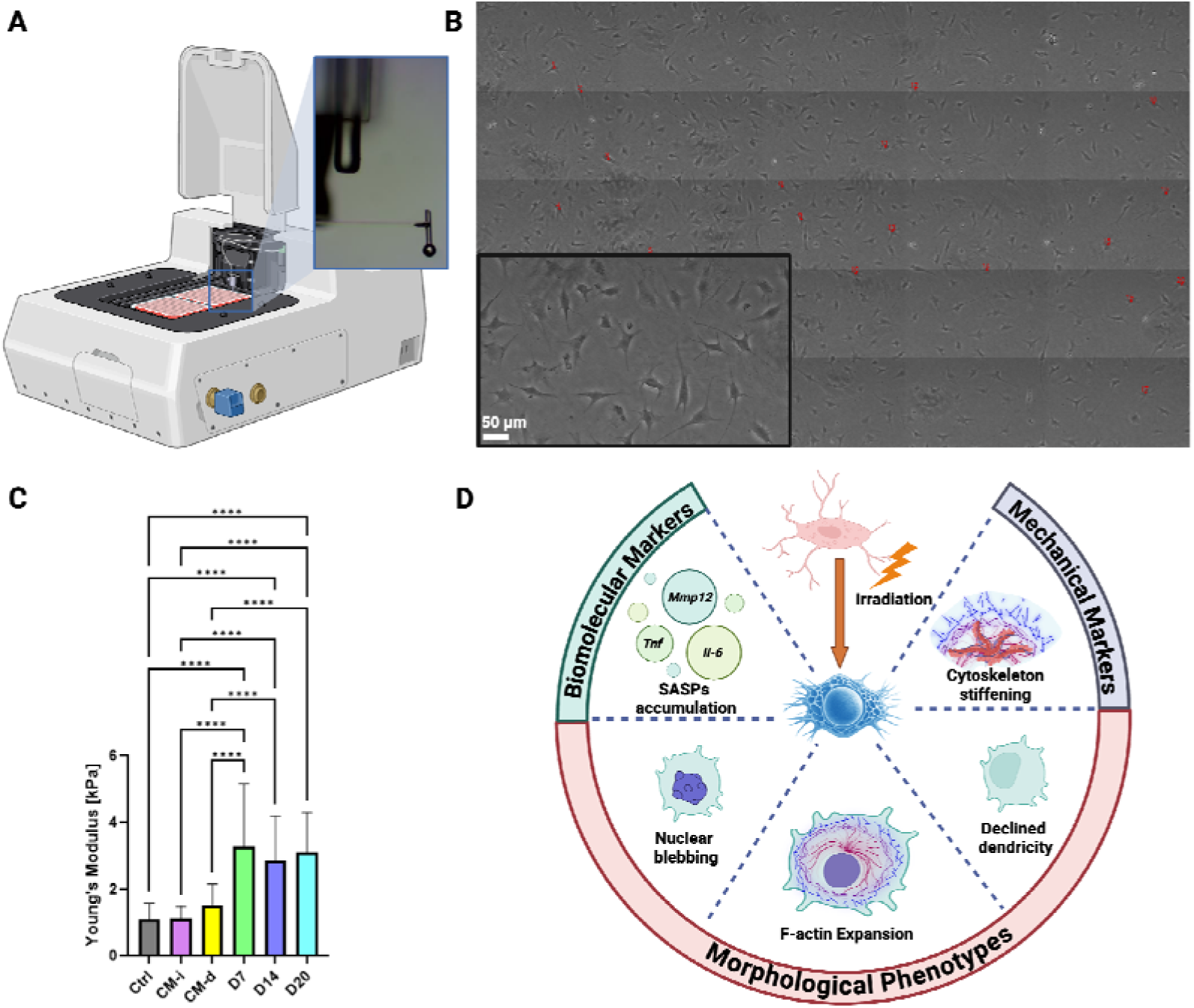
Irradiated osteocytes exhibit significant increases in cytoskeletal stiffness. **A)** Representative image of our live single-cell mechanical testing setup. **B)** Live cells attached to the bottom of the cell culture plate being indented. Red dots indicate the probe size and locations of indentation. Th main image was obtained via image stitching, and the inset magnifies an individual indentation site for better visibility. **C)** Young’s moduli of osteocytes cytoskeleton measured using Hertzian contact mechanic model. Data from 50 cells per condition are presented as mean ± SD. Significance levels are denoted as *: p < 0.05, **: p < 0.01, ***: p < 0.001, ****: p < 0.0001. **D)** Summary of the study: Irradiation-induced senescence in osteocytes is characterized by changes in micromechanical properties, morphological features, and biomolecular markers.

The stiffening of the osteocyte cytoskeleton, coupled with the observed senescence-associated heterogeneity in cytoskeletal organization (**Figure 4A**), suggests a decline in cell motility and possibly mechanosensory properties. Such degenerative changes potentially lead to deterioration in interactions of osteocytes with the ECM, a phenomenon similarly reported in studies involving fibroblasts.[88] The influence of substrate stiffness on cellular behavior, particularly the SASP secretion in senescent cells, has been previously studied in other cell types.[89] Gresham et al. demonstrated that compliant substrates (e.g., polyacrylamide hydrogels with a stiffness of 50 kPa) can mitigate SASP secretion in stress-induced senescent MSCs. [89] This reduction was evident in key pro-inflammatory markers and DNA damage-associated signals (e.g., γH2AX). Their findings highlight the importance of substrate mechanical properties in regulating senescence-associated phenotypes. In our study, all experimental groups were cultured on the same rigid tissue culture plastic (TCP, ∼1 GPa) coated with rat-derived collagen type I, ensuring that substrate effects were applied uniformly across all groups. This uniformity minimizes substrate stiffness as a confounding factor in our comparative analyses of cytoskeletal stiffening and SASP-related alterations between irradiated and control osteocytes. However, it is noteworthy that the rigid substrate likely exerted greater mechanical resistance compared to compliant hydrogels or native bone matrix, which may have amplified cytoskeletal stiffening and SASP-related changes in our study. These findings underscore the importance of exploring compliant and tunable stiffness substrates in future studies to better replicate the mechanical properties of the 3D bone microenvironment and refine our understanding of senescence in osteocytes.

In vivo, osteocytes are embedded within the mineralized bone matrix, which exhibits a highly heterogeneous and dynamic mechanical environment.[23,90,91] Cortical bone has a stiffness range of 17–20 GPa, while trabecular bone exhibits stiffness values between 10 and 3,000 MPa, depending on its density and microarchitecture.[90,91] This mechanical environment is critical for osteocyte mechanosensation, as mechanical forces are transmitted through the extracellular matrix and dendritic networks to regulate bone remodeling.[23] The high stiffness of the in vitro TCP substrate differs markedly from the in vivo conditions, potentially amplifying stress-induced phenotypes such as cytoskeletal stiffening and altered mechanotransduction.

These findings indicate that while irradiated osteocytes consistently exhibit cytoskeletal stiffening, systemic exposure to SASP factors results in less significant changes in cytoskeletal stiffness. Future studies employing 3D culture systems with tunable stiffness substrates or more physiologically relevant bone-mimetic scaffolds are needed to validate these findings and assess the role of substrate mechanical properties in osteocyte mechanosensory function.

## 4. Conclusions and Future Directions

This longitudinal study highlights the impact of continuous local SASP factor accumulation, resulting from irradiation, on distinct senescence-associated biophysical markers, supported by corresponding biomolecular changes (**Figure 5D**). Importantly, senescent osteocytes in this study exhibit SASP profiles consistent with those observed in other cell types, including increased secretion of pro-inflammatory cytokines such as *Il-6* and matrix-degrading enzymes such as *Mmp-12* (**Figure 1**).[92] Our findings demonstrate that local SASP accumulation has a pronounced effect on osteocyte cytoskeletal properties, including significant stiffening and heterogeneity in cytoskeletal organization. To build on these findings, future research could incorporate Western blot analysis to validate the observed senescence-associated changes at the protein level. This approach would complement and support the RT-qPCR data by confirming that transcriptional changes in key markers, such as *p16* and *p21*, are reflected at the protein level. Additionally, flow cytometry (FACS) could provide direct evidence of cell cycle disruption by analyzing the distribution of cells across different phases of the cell cycle. Additionally, future studies will expand upon this work by investigating changes in receptors for SASP cues and quantifying the bioactivity of secreted SASP proteins. These analyses will provide deeper insights into the mechanisms through which SASP factors mediate their effects and their functional consequences on cellular and tissue-level remodeling. Such efforts will further inform the development of targeted therapeutic strategies to mitigate the deleterious effects of cellular senescence.

The observed increase in cytoskeletal stiffness in senescent osteocytes also aligns with previous studies in senescent fibroblasts, which exhibit alterations in the cytoskeleton and changes in actin organization leading to altered cell morphology and diminished proliferation and migration.[86] Similarly, studies in MSCs have shown that irradiation-induced senescence leads to altered cytoskeletal and focal adhesion organization and reduced cellular migratory capacity.[93] These parallels highlight the conserved nature of cytoskeletal alterations across different cell types and expand our understanding of osteocyte responses to localized and systemic senescence effects. Notably, in our present study, systemic SASP exposure (CM groups) did not induce substantial changes in cytoskeletal stiffness or morphological properties compared to controls, suggesting a more localized influence of SASP on osteocyte biophysical properties. However, trends observed in the CM-d group point to the potential role of cellular maturation in modulating SASP uptake, warranting further investigation.

Osteocyte dysfunction, driven by senescence-associated cytoskeletal changes, plays a pivotal role in the pathophysiology of osteoporosis.[5,92] Cytoskeletal dynamics are integral to osteocyte morphology and function, enabling these cells to sense and transduce mechanical signals via their dendritic processes.[23] The observed cytoskeletal stiffening and altered organization likely impair mechanosensing, reducing the ability of osteocytes to adapt to mechanical stimuli and coordinate bone remodeling.[94] Furthermore, SASP, characterized by inflammatory cytokines and matrix-degrading enzymes, exacerbates bone fragility by promoting osteoclast activity and inhibiting osteoblast function,[92,95] processes involved in bone remodeling and tightly regulated by osteocytes. This interplay between cytoskeletal changes and SASP secretion highlights the potential contribution of senescent osteocytes to the microarchitectural deterioration observed in osteoporosis, a condition closely linked to age-related imbalance in bone remodeling, reduction in osteocyte viability/function, and chronic inflammation that contributes to progressive bone loss and increased fragility over time.[96]

While these findings provide a robust framework for understanding osteocyte dysfunction, the use of rigid tissue culture plastic (TCP, ∼1 GPa) in our study may have amplified stress-induced phenotypes. In vivo, osteocytes are embedded within a highly heterogeneous bone matrix, with stiffness values ranging from 17–20 GPa in cortical bone to 10–3,000 MPa in trabecular bone, depending on density and architecture.[90,91] Furthermore, 2D cultures may not entirely capture the intercellular interactions and extracellular matrix dynamics found in living tissues, which are crucial for cellular behavior and function. The pronounced differences between in vitro and in vivo mechanical environments underscore the importance of future studies using 3D culture systems or bone-mimetic scaffolds that better replicate physiologically and clinically relevant conditions. Another limitation of this study was the inability to characterize the same senescent cells across multiple assays, which would establish direct correlations between senescence markers and functional outcomes. Single-cell approaches, such as RNA sequencing or multi- modal profiling, could address this limitation by revealing heterogeneity within senescent cell populations and uncovering relationships between SASP profiles and biophysical changes. However, these methods are resource-intensive and currently impractical for large-scale studies.

Future research integrating such advanced assays could provide deeper insights into the dynamic and heterogeneous nature of senescence, particularly in osteocytes, and their role in bone remodeling.

To address these limitations, future studies will investigate the relationship between cytoskeletal stiffening, motility, and mechanosensory properties in osteocytes using live-cell imaging of dendritic processes and 3D culture models with tunable stiffness. Additionally, combining our nanoindentation system with real-time PCR or immunocytochemistry will allow us to explore key mechanosignaling pathways, such as YAP/TAZ and FAK, to elucidate the molecular mechanisms underlying these changes. The inclusion of proteomic measures in future studies could enhance the characterization of SASP factors and offer a more comprehensive understanding of senescence processes. By establishing a causal link between cytoskeletal stiffening, mechanotransduction impairment, and osteocyte-driven bone remodeling dysfunction, these efforts aim to validate our findings and uncover potential therapeutic targets for mitigating prevalent age-related bone diseases like osteoporosis.

## Author Contributions

MT designed the study, experiments, and concept. JL and MT performed the experiments, researched, analyzed, interpreted the data, and wrote the original manuscript. CK, HS, MAP, and MT analyzed, interpreted the data, and revised manuscript. MT, JLK, and KMM provided resources. All authors reviewed and revised the manuscript. All authors read and approved the final manuscript.

## Acknowledgements

This research and MT are supported by the Walker Department of Mechanical Engineering at The University of Texas at Austin. KMM is supported by NCI (RO1 CA198279 and CA250905).

## Data Availability

The data that support the findings of this study are available on request from the corresponding author.

## Supplementary Materials

**Figure S1:**
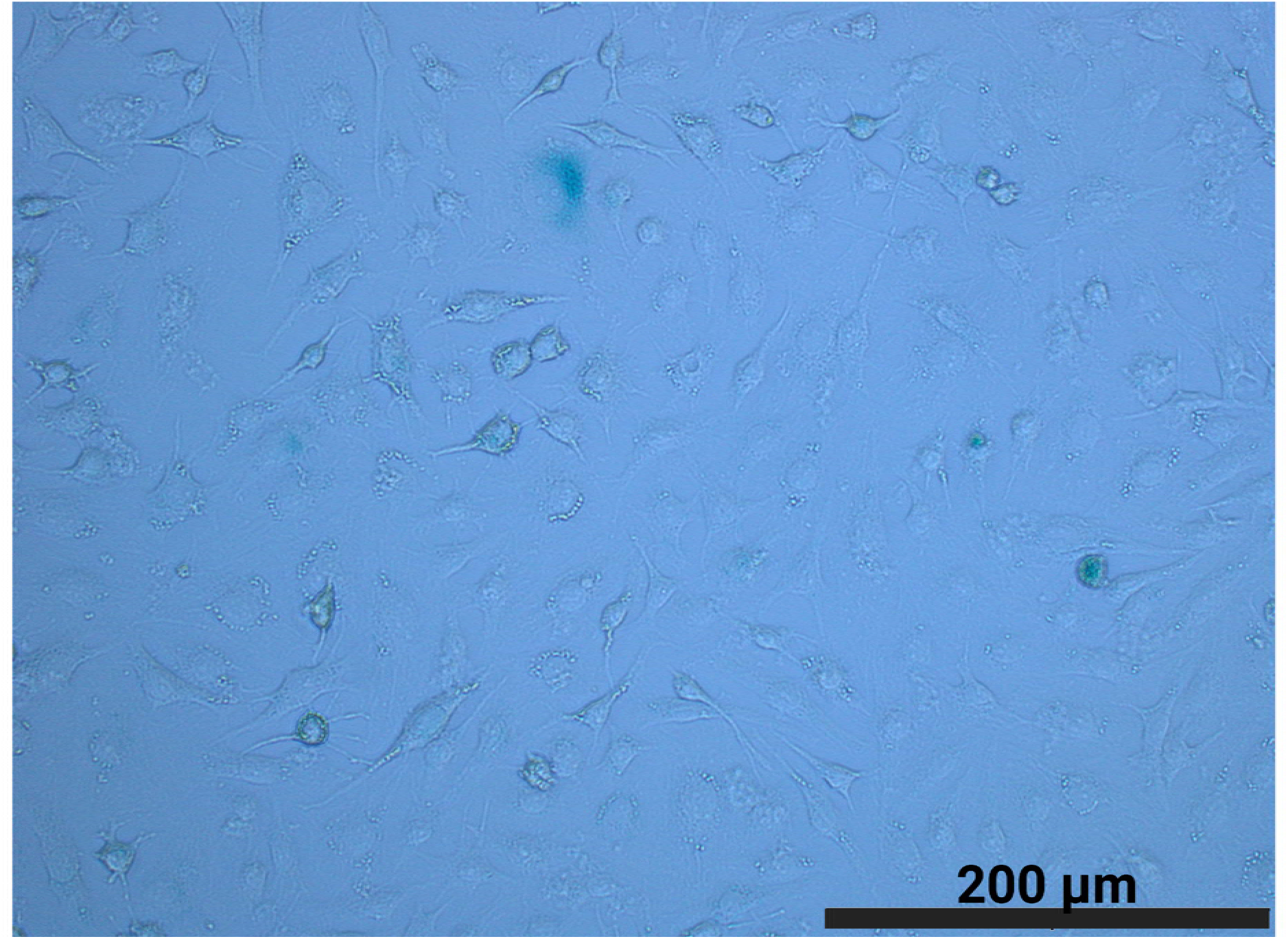
Representative SA-β-Gal staining images demonstrating senescence in the CM-d-treated osteocytes.

**Figure S2:**
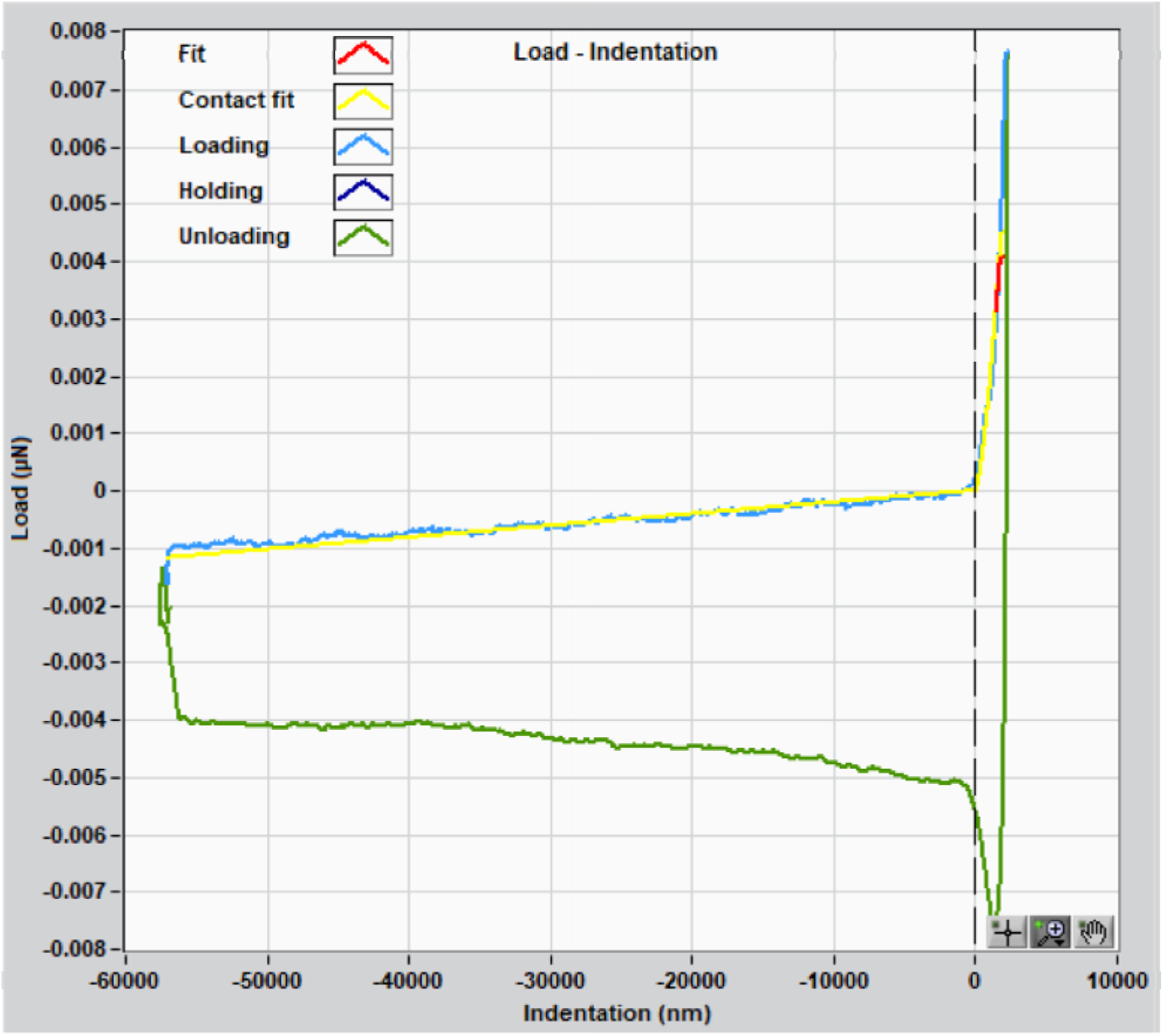
Nanoindentation curve fitting over the linear region demonstrates the application of Hertzian contact mechanics to determine elastic modulus values of osteocytes.

**Figure S3:**
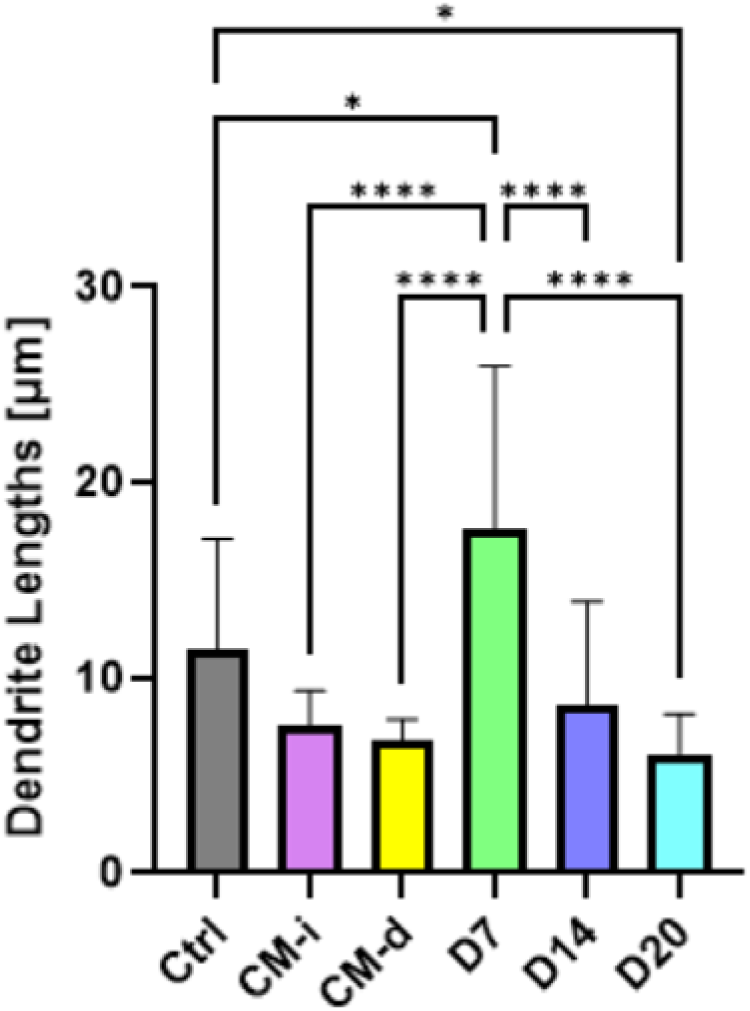
Quantitative analysis of average dendrite lengths in osteocytes under different experimental conditions. Sample size for each group is n = 3, presented as mean ± SD. *: p < 0.05, **: p < 0.01, ***: p < 0.001, ****: p < 0.0001.

**Figure S4:**
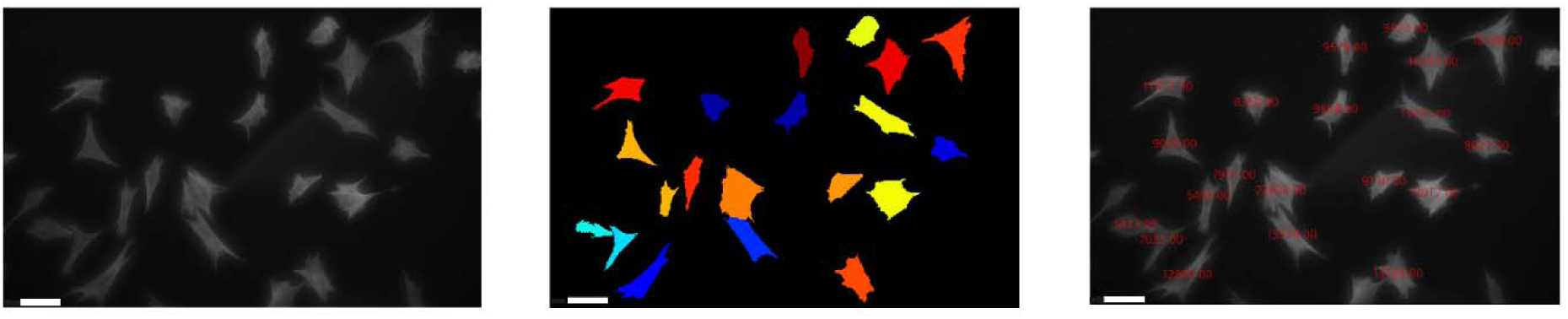
Example of segmentation (center) and quantification (right) of F-actin micrographs (left) obtained via IF staining, analyzed using CellProfiler. Scale bar: 50 µm. Units in the quantification image (right): pixel area.

**Figure S5:**
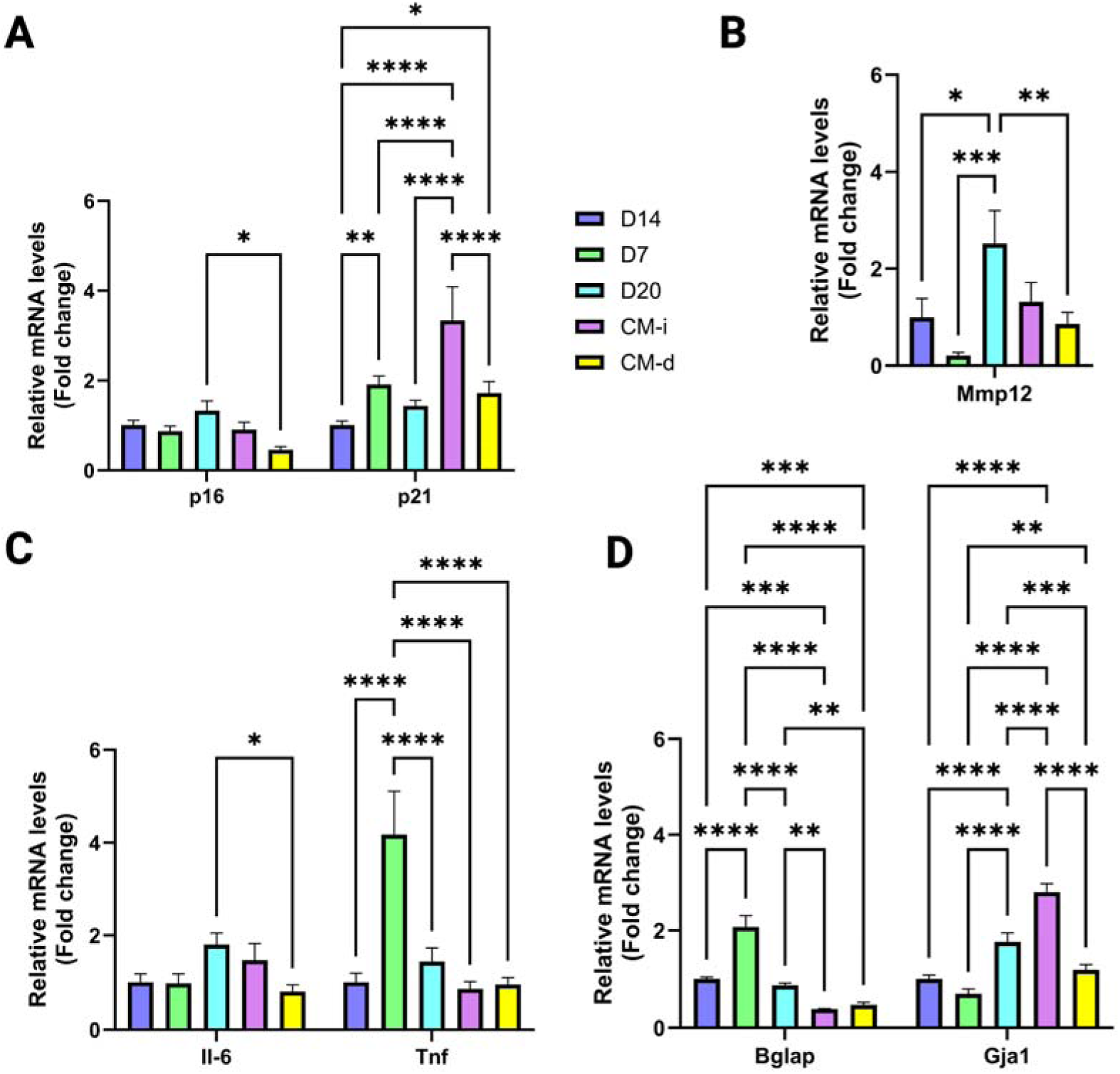
**A)** RT-qPCR results of senescence, **B-C)** SASP factors for all conditions, and **D)** osteocyte markers, normalized to *Gapdh* housekeeping gene expression and compared to the Day 14 post-irradiation group. n = 3, mean ± SD. *: p < 0.05, **: p < 0.01, ***: p < 0.001, ****: p < 0.0001.

**Table S1:**
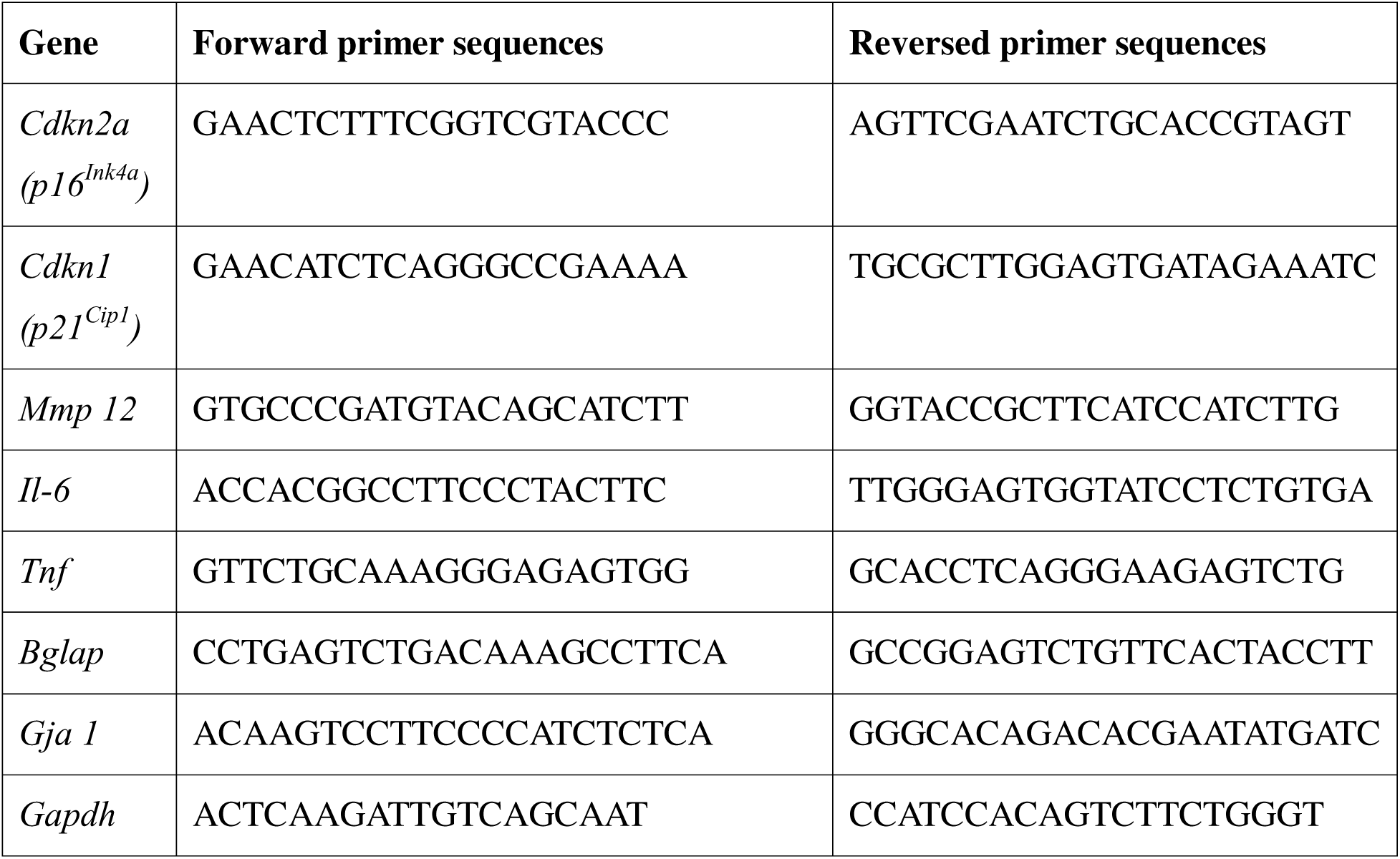
Primer sequences used for RT-qPCR experiments.

